# Cannabidiol improves learning and memory deficits and alleviates anxiety in 12-month-old SAMP8 Mice

**DOI:** 10.1101/2023.12.21.572902

**Authors:** Monica N. Goodland, Subhashis Banerjee, Michael L. Niehoff, Benjamin J. Young, Heather Macarthur, Andrew A. Butler, John E. Morley, Susan A. Farr

## Abstract

Cannabidiol (CBD) has gained a lot of interest in recent years for its purported medicinal properties. CBD has been investigated for the treatment of anxiety, depression, epilepsy, neuroinflammation, and pain. Recently there has been an interest in CBD as a possible treatment for age-related disorders such as Alzheimer’s disease and related disorders (ADRD). Here we tested the hypothesis that chronic CBD administration would improve learning and memory in the SAMP8 mouse model of Alzheimer’s disease. SAMP8 mice aged 11 months (at the start of the study) were administered vehicle or CBD (3 or 30 mg/Kg) daily via oral gavage for 2 months. Vehicle-treated young SAMP8 mice (age 3 months at the start of the study) served as unimpaired controls. After 30 days of treatment (4 and 12 months of age), learning and memory, activity, anxiety, strength and dexterity were assessed. High dose CBD treatment significantly improved learning and memory of the 12-month-old mice in the T maze. Novel object recognition memory was also improved by CBD in aged CBD treated mice. Aged CBD treated mice also displayed less anxiety in the elevated plus maze test compared to controls. However, activity and strength levels were similar between groups. Biochemical analysis revealed decreased markers of oxidative stress, providing a possible mechanism by which CBD treatment impacts learning, memory, and anxiety. These results highlight the potential use of CBD as a therapeutic for age related cognitive impairment and dementia.

## INTRODUCTION

Cannabidiol (CBD) is an abundant phytocannabinoid found in the *Cannabis sativa* plant (1). While CBD is not noneuphorigenic and does not mediate the responses observe with the consumption of cannabis (2), it is being investigated as a pharmacological agent for treating a variety of neurological conditions. CBD is being investigated for the treatment of spasticity associated with multiple sclerosis, as an anxiolytic, as an antiemetic, and for treating chronic pain (3-6). One drug containing a highly purified form of CBD (Epidioloex) has been approved by the FDA for treating severe childhood seizures observed in Lennox - Gastaut syndrome (LGS), Dravet syndrome (DS), or tuberous sclerosis complex (TSC) (7). CBD has a diverse range of molecular targets including GPCRs and ion channels. CBD is an agonist at the 5-HT_1A_ and TRPV1 receptors where it has anxiolytic and analgesic effects (8) as well as a partial agonist for the D2 receptor. CBD also works at G protein coupled receptor 55, CB1 and CB2 as an antagonist and a partial (9-11). (12). Cannabinoids also regulate gene expression by acting as ligands for peroxisome proliferator activated receptors (PPARs) (13) (See de Almeida and Levi, 2020 (8) for a complete review).

Recently, there has been a focus on whether CBD could have a therapeutic role in treatment of age-related disorders such as cognitive, mood, and muscle disorders (4, 14, 15). CBD has been known to inhibit Aβ induce neuro inflammation one of the neuropathologies that underlie the development of Alzheimer’s disease (AD)(16)., progressive neurodegenerative disorder that causes cognitive impairment, culminating in dementia and eventually death (17). The single greatest risk factor for the development of AD is age, with 1 in 10 people aged 65 years or older carrying the diagnosis (18). Currently, there are few medications approved for the treatment of cognitive impairment caused by AD, and they all have limited efficacy for patients (19). CBD has been investigated as a potential treatment due to its antioxidant and anti-inflammatory effects (20).

Here, we examine the potential use of CBD in treating age-related memory loss, anxiety, strength and dexterity impairment in the Senescence-accelerated mouse -prone 8(SAMP8) mice, a polygenic model of spontaneous onset AD (21). The mice develop early onset age-related deficits in learning and memory with impairment in learning and memory starting at about 8 months of age and being severe by 12 months of age (22, 23). The memory deficits correspond to age-related increases in amyloid beta protein (Aβ), hyperphosphorylated tau, oxidative stress, neuroinflammation and impaired efflux of Aβ across the blood-brain barrier (24-28). The SAMP8 also has accelerated loss of skeletal muscle (29). Eleven-month-old mice at the start of treatment were given CBD and compared to a young 3-month-old SAMP8 mouse (at the start of treatment) to determine if it could reverse their age-related memory deficits.

## Materials and Methods

### Animals

Young (3 mos of age) and aged (11 mos of age) male SAMP8 mice were obtained from our breeding colony. A young and an aged mice received only vehicle (medium chain triglycerides (MCT). The other 2 groups of aged mice received either 3 or 30 mg/Kg /day CBD respectively for 8 weeks. These 2 doses were chosen based on dosages of CBD recommended to consumers by BeLeaf. Compounds were administered by oral gavage 75ul/mouse. All studies were conducted with the approval of the Animal Care and Use Committee at Saint Louis University. Sentinel animals are tested regularly to ensure the facility is pathogen free. Food and water were available on an *ad libitum* basis and the vivarium had a 12 h light-dark cycle with lights on at 0600 h. Behavioral experiments were conducted between 0730 and 1430.

### CBD

The full spectrum CBD compound and vehicle (MCT) were supplied by BeLeaf (Earth City. MO). The oil is extracted via an ethanol extraction method. Dry plant material is soaked in 200 proof organic ethanol for 1 hour. It is then put in a traditional RotoVap to evaporate the alcohol off the oil. To remove any residual alcohol, it is put in a vacuum oven at 90 degrees for 12 hours. What is left is the pure crude CBD oil with the majority of everything that is in the plant material. The extra concentration potency was 81.70% CBD, independently verified by LK Pure Lab (Sparta, IL) (See supplement for entire breakdown of the extract). Mice were weighed at commencement of the study, and then weekly to accurately prepare dosages based on average group body weight. Mice were treated for 8 weeks. Behavioral testing began after 4 weeks of treatment.

### Cognitive testing

Behavioral testing began during the fifth week of treatment when the younger mice were 4 mos old and the older mice were 12 mos old. Novel object recognition (NOR) is a declarative memory task that involves the hippocampus (30) when the retention interval is 24 hours after the initial exposure to the objects. Mice were habituated to the empty apparatus for 5 minutes a day for 3 days prior to introduction to objects. During the training session, mice were exposed to two similar objects and allowed to explore for 5 minutes while the observer recorded time spent investigating each object. The apparatus and objects were cleaned between each mouse. Twenty-four hours later, mice were exposed to one of the original objects and a novel object. The time spent investigating each object was again recorded. The novel object was made from the same material as the original object and approximately the same size but had a different shape. The test is based on the tendency of mice to spend more time exploring new objects than familiar ones. Thus, the more time spent with the novel object, the greater the retention/memory of the familiar object at 24 hours, as defined by the discrimination index (DI). DI= (time spent with novel object – time spent with familiar object)/(total time spent with both objects).

The aversive T maze is both a learning task based on working memory and a reference memory task (31). The T maze consisted of a black plastic alley with a start box at one end and two goal boxes at the other. The start box was separated from the alley by a plastic guillotine door that prevented movement down the alley until raised during training trials. An electrifiable floor of stainless-steel rods ran throughout the maze to deliver a mild, scrambled foot-shock. Mice were not permitted to explore the maze prior to training. A block of training trials began when a mouse was placed into the start box. The guillotine door was raised, and a cue buzzer sounded simultaneously; 5 seconds later foot-shock was applied. The arm of the maze entered on the first trial was designated “incorrect” and the mild foot shock was continued until the mouse entered the other goal box, which in all subsequent trials was designated as “correct” for that mouse. At the end of each trial, the mouse was returned to its home cage until the next trial. Each mouse was trained until he made one-avoidance. Training used an inter-trial interval of 35 seconds, the buzzer was of the doorbell type sounded at 55 dB, and shock was set at 0.35 mA (Coulbourn Instruments scrambled grid floor shocker model E13-08). Retention was tested one week later by continuing training until mice reached the criterion of 5 avoidances in 6 consecutive trials. The results were reported as the number of trials to reach criterion for the retention test.

To eliminate changes in locomotor activity and anxiety as a factor affecting performance on subsequent behavioral tests, mice subjected to an open field test. Mice were placed in an open field (67 cm x 57 cm x 24 cm) for 5 minutes. Distance traveled, time spent in center and peripheral zones, and number of entries into the center zone (33.5 cm x 28.5 cm) of the open field was recorded by an ANYmaze tracker (San Diego Instruments, San Diego, CA).

To further test for anxiety as a factor affecting performance behavioral tests, mice were subjected to an elevated plus maze test. Mice were placed onto the center zone of an elevated plus maze apparatus consisting of 4 perpendicular arms in the shape of a plus sign, elevated 50 cm above the floor. Each arm is 35.5 cm in length; two opposite arms are enclosed and two are open. The test consisted of placing a mouse in the central platform facing an enclosed arm and allowing it to freely explore for 5 minutes. The test arena was wiped with a damp cloth after each trial. The number of entries into the open and closed arms and the time spent in the open arms was recorded by an ANYmaze tracker (San Diego Instruments, San Diego, CA). Anxiolysis is indicated by increased time spent in open arms and number of open arm entries.

### Strength and Dexterity

Since CBD has been found to improve muscle function (14), we examine the effects of chronic treatment on the age-related decrease in strength seen in a separate set of SAMP8 (n=6∼8 per group).

Grip Strength Testing the animal’s front paws are placed on a wire grid, which the animal will naturally hold on to while its tail is gently pulled backwards. The maximum strength of the grip prior to grip release is recorded using a grip Strength meter (Model #05123, Columbus Instruments, Columbus, OH). Values were corrected by dividing the values detected by the strength meter by body weight in kilograms.

Cage hang the mouse was allowed to grip the top of a wire mesh cage, which was then inverted 12 inches over a 3/4 inch foam pad. The time the mouse was able to hang was recorded, the test ending after a maximum of 3 min.

Rod test a mouse was placed on a pole 1/4 inch in diameter 12 inches over a 3/4 inch foam and the time the mouse balanced on the rod recorded. The test was stopped after 3 min.

String hang a mouse gripped a string suspended 12 inches above a 3/4 inch foam pad. The time that a mouse was able to hang on the string was recorded with a maximum hang time of 3 min.

### RNA extraction, cDNA synthesis and real-time quantitative RT_qPCR

Brains were collected and flash frozen. Rt-qPCR was run on cortex tissue to examine changes in genes related to oxidative stress. We have previously shown that the SAMP8 are an excellent model to study oxidative changes related to aging (32). The extraction of the total RNA from was performed by using kits from Ambion® RiboPure™ (AM1924, ThermoFisher Scientific, Waltham, MA,. cDNA was synthesized with cDNA reverse transcription kit from Quanta. PCR was conducted using a QuantStudio Realtime PCR machine, Applied Systems using primers from Integrated DNA Technology. Data were normalized using three reference genes (HPRT1, 36B4, PPIB) (See Supplement 2 for list of gene primer sequences).

### Immunoblotting

Protein was isolated by standard protein lysis RIPA buffer, containing the protease-phosphatase inhibitor tablet (A32965, Thermo-Scientific, Waltham, MA), and quantified using a BCA protein assay kit (23225, Pierce; ThermoFisher Scientific, Waltham, MA). Immunoblotting was done following the standard procedure (33). An equal amount of protein was loaded and blotted for 4-HNE (MA527570, Life Technologies Corpa, Carlsbad, CA), Anti-NOX/gp91phox (ab106940, abcam, Cambridge, MA) and HSP90 (4874 Cell Signaling Technology, Danvers, MA) using antibodies..

### Statistical analysis

Results were analyzed using analysis of variance (ANOVA) to examine the effect across groups. The measure of acquisition and retention in the T-maze were the number of trials to reach criterion. The results for object recognition are presented discrimination index based on total time with the novel object out of total exploration time. Biochemistry was analyzed using a one-way ANOVA followed by a Tukey’s post hoc test. Results are expressed as means with their standard errors. Tukey’s or Dunnett’s post hoc analysis was used to compare means between groups. Outliers defined as values that were more than two standard deviations from the group mean were removed from the analysis.

## Results

### T-maze

We examined the effect of CBD on T-maze foot shock avoidance and novel objection recognition. An ANOVA on trials to first avoidance during acquisition was significant F(3,42) = 118.83, P<0.0001. Tukey’s post hoc test indicated that the 12 mos-old SAMP8 mice receiving vehicle took significantly more trial to make one avoidance during acquisition than the 4 mos-old SAMP8 mice receiving vehicle P<0.0001. Tukey’s post hoc test indicated that the 12 mos-old SAMP8 mice receiving 3 and 30 mg/Kg of CBD mice took significantly fewer trials to make one avoidance than the 12-mos-old SAMP8 mice that received vehicle P<0.05 and P<0.0001. The mice that received the 3 m/Kg took significantly longer to make one avoidance than the 4 mos-old mice receiving vehicle. There was no difference between the 4 mos-old SAMP8 mice that received vehicle and the 12 mos-old mice that received 30mg/Kg of CBD (Figure 1A). The ANOVA for latency to escape the shock on trial one of acquisition was not significant F(3,42) = 0.1201, P NS (Data not shown). The ANOVA for trials to criterion for the retention test was significant F(3,42) = 7.32, P<0.0005. Tukey’s post hoc test indicated that the 12 mos-old SAMP8 mice that received vehicle took significantly longer to make criterion of 5 avoidance in 6 consecutive trials than the 4 mos-old SAMP8 mice that received vehicle P<0.001. The mice that received 30 mg/Kg of CBD took significantly fewer trials than the 12mos-old SAMP8 mice that received vehicle P<0.05. There was no difference between the 12 mos-old mice that received 3mg/Kg CBD and any of the other groups (Figure 1C).

**Figure 1.**
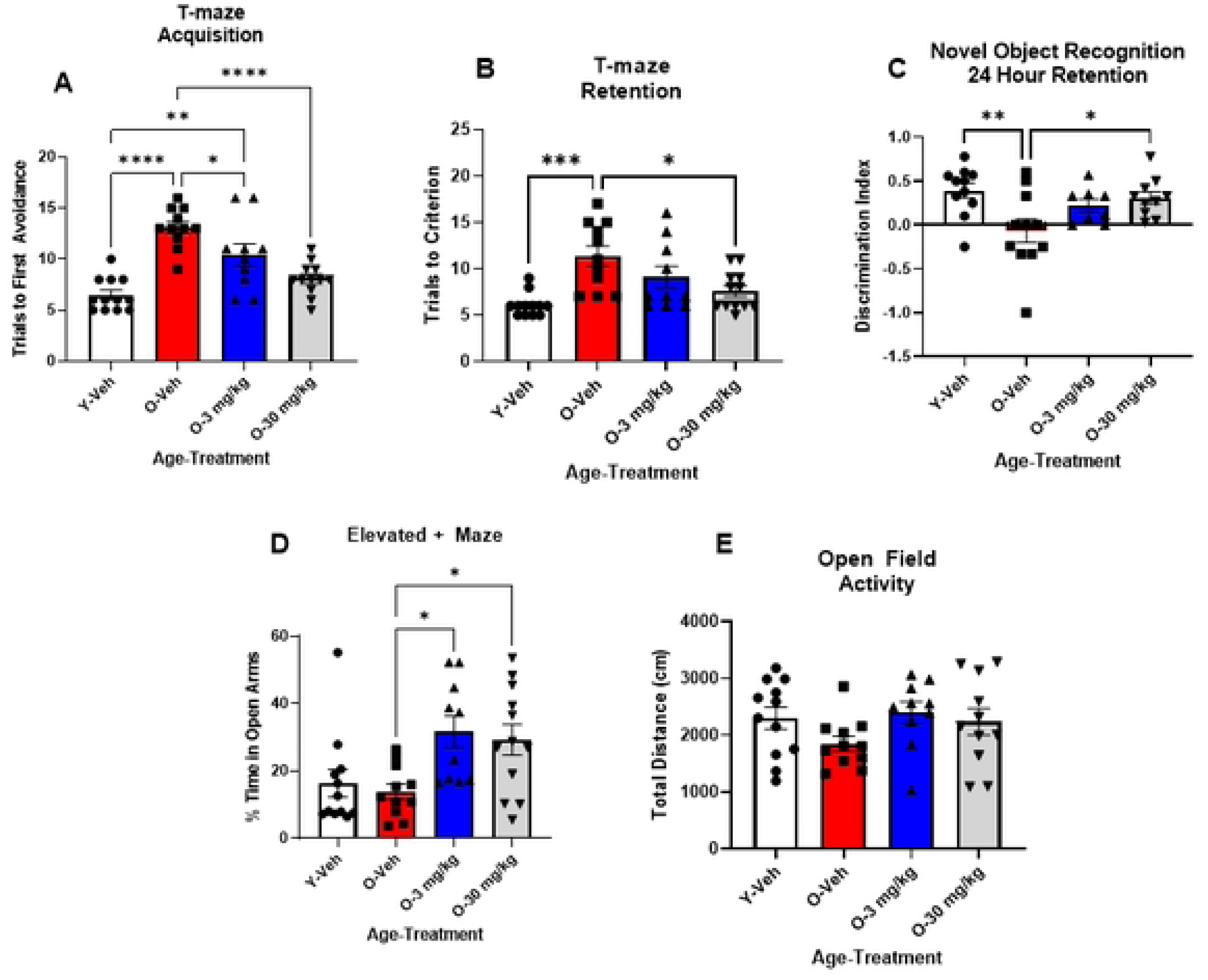
CBD treatment improves deficits of learning and memory in aged male SAMP8 mice and is anxiolytic. **A**: Vehicle treated aged (12 mos-old) male mice (red bar) showed deficits in memory and learning in the T-maze compared to vehicle treated young (4 mos-old) male mice (white bar). 12 mos-old SAMP8 mice that received CBD 30 mg/Kg/daily (gray bar) (O-30mg/Kg) showed improvement in learning and memory compared to vehicle treated aged mice (red bar), whereas aged mice that received 3 mg/Kg (blue bar) (O-3mg/Kg) showed only improvement in learning, but not memory (N=10∼12 per group). **B**: In NOR with a 24- hour retention interval aged (12 mos old) mice receiving vehicle (red bar) had significantly reduced memory of a previously seen object compared to young (4 mos-old) vehicle treated mice (white bar). 12 mos-old mice that received CBD 30 mg/Kg (gray bar) but not 3 mg/Kg (blue bar) (O-30mg/Kg) showed significant improvement in memory of the previously seen object, compared to the vehicle treated 12 mos-old mice (N=8∼11 per group) **D**: The elevated maze test showed that 12 mos-old mice (red bar) and 4 mos-old mice (white bar) that received vehicle had similar anxiety levels to one another. 12 mos-old mice that received both 3 (blue bar) or 30 mg/Kg (gray bar) of CBD showed decreased anxiety compared to the 12 mos-old mice that received vehicle (red bar) (O-Veh) (N=10∼12 per group). **E:** There were no differences in general activity between any of the groups of mice in an open field test (N=10∼12 per group). *P<0.005, **P<0.01, *** P<0.001, ****P<0.0001.

### Novel Object Recognition

The ANOVA for the discrimination index at 24 hour retention test was significant F(3,36) = 4.399, P<0.01. Tukey’s post hoc test indicated that the 12 mos-old SAMP8 mice had a significantly lower discrimination index indicating they spent less time with the novel object P<0.01. The 12 mos-old SAMP8 mice that received 30 mg/Kg of CBD spent significantly greater time with the novel object than the 12 mos-old SAMP8 mice that received vehicle P<0.05. There was no difference between the 12 mos SAMP8 group that received 3 mg/Kg CBD and any of the other groups (Figure 1C). The ANOVA for exploration of the like objects on day 1 of novel object recognition was not significant F(3,36) = 2.35, P NS (Data not shown).

### Elevated plus maze

The ANOVA for time spent in the open arms of the elevated + maze was significant F(3,42) = 6.427, P<0.001. Tukey’s post hoc test indicated that the 12 mos-okl SAMP8 mice that received 3mg/Kg or 30mg/Kg CBD spent significantly more time in the open arms than the 12 mos-old SAMP8 that received vehicle (P<0.05), but not compared to the 4 mos mice that receive vehicle. There were no differences between the 4 mos-old SAMP8 and the 12 mos-old SAMP8 mice that received vehicle (Figure 1D).

### Open Field Activity

The ANOVA for distance traveled in an open field was not significant F(3,42) = 1.354, P NS (Figure 1E).

### Grip Strength

The ANOVA for forearm grip strength was significant F(2, 24) = 4.95, P<0.008. Tukey’s post hoc test indicated that the 4 mos SAMP8 mice had significantly greater grip strength than the 12 mos-old mice that received vehicle and the 12 mos old mice that received 30 mg/Kg of CBD. (Figure 2A).

**Figure 2.**
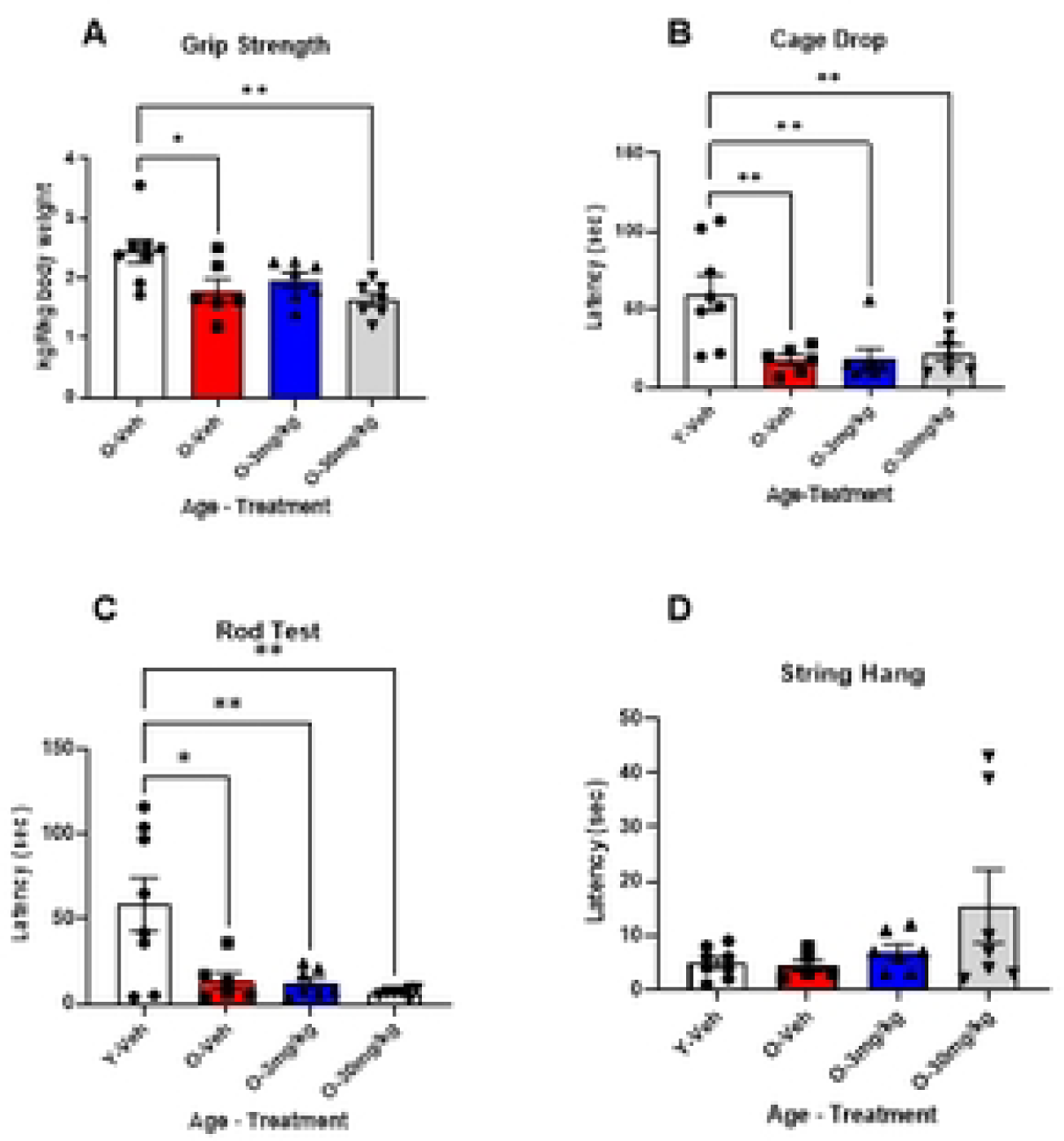
CBD treatment did not improve aging-related declines in strength of SAMP8 mice. The young 4-mos-old vehicle treated SAMP8 mice (white bars) were significantly stronger in 3 measures of strength: **A,** grip meter, **B**, cage drop and **C**,rod test compared to the 12 mos- old vehicle treated SAMP8 mice (red bars). Neither 3 (blue bars) nor 30 mg/Kg (gray bars) CBD treatment had any effect on these measures. **D**: There were no effects of either age, or treatment, in the string hang test. (N = 6-8 per group for all) * P<0.05, ** P<0.01.

### Cage Drop

The ANOVA for cage drop was significant F(3, 24) = 7.323, P<0.001. Tukey’s post hoc analysis indicated that the 4 mos- old SAMP8mice were able to hang upside down by all 4 paws for significantly longer than any of the 12 mos old groups (Figure 2B). There was no difference between the 12 mos-old groups.

### Rod Test

The ANOVA for the Rod Test was significant F(3, 24) = 7.25, P<0.0001. Tukey’s post hoc test indicated that the 4 mosoldSAMP8 mice stayed on the rod significantly longer than any of the 12 mos old mice (Figure 2C).

### String Hang

The ANOVA for forearm string hang was not significant (3,23) = 1.96, P NS (Figure 2D).

### CBD treatment improves RNA and protein markers of oxidative stress

qPCR analysis of markers of oxidative stress indicated that SOD2, SOD1, and TXNRD1 had significantly higher expression in the 12 mos vehicle treated SAMP8 mice compared to the 4 mos vehicle treated SAMP8 mice P<0.01 and CAT, GSTM2 and PRDX3 were significantly elevated in the 12 mos vehicle treated mice compared to the 4 mos vehicle treated (P<0.05). CBD at 30 mg/Kg was able to significantly reduce SOD2 and SOD1,P<0.05 (Figures 3A, B,C, F, H & I) while CAT, TXNRD1, GSTM2 and PRDX3 were not significantly reduced with 3mg/Kg or 30 mg/Kg CBD treatment, they were no longer significantly elevated compared to the 4-mos SAMP8 mice suggesting an overall reduction in oxidative stress (Figures 3 C, F, H and I).

**Figure 3.**
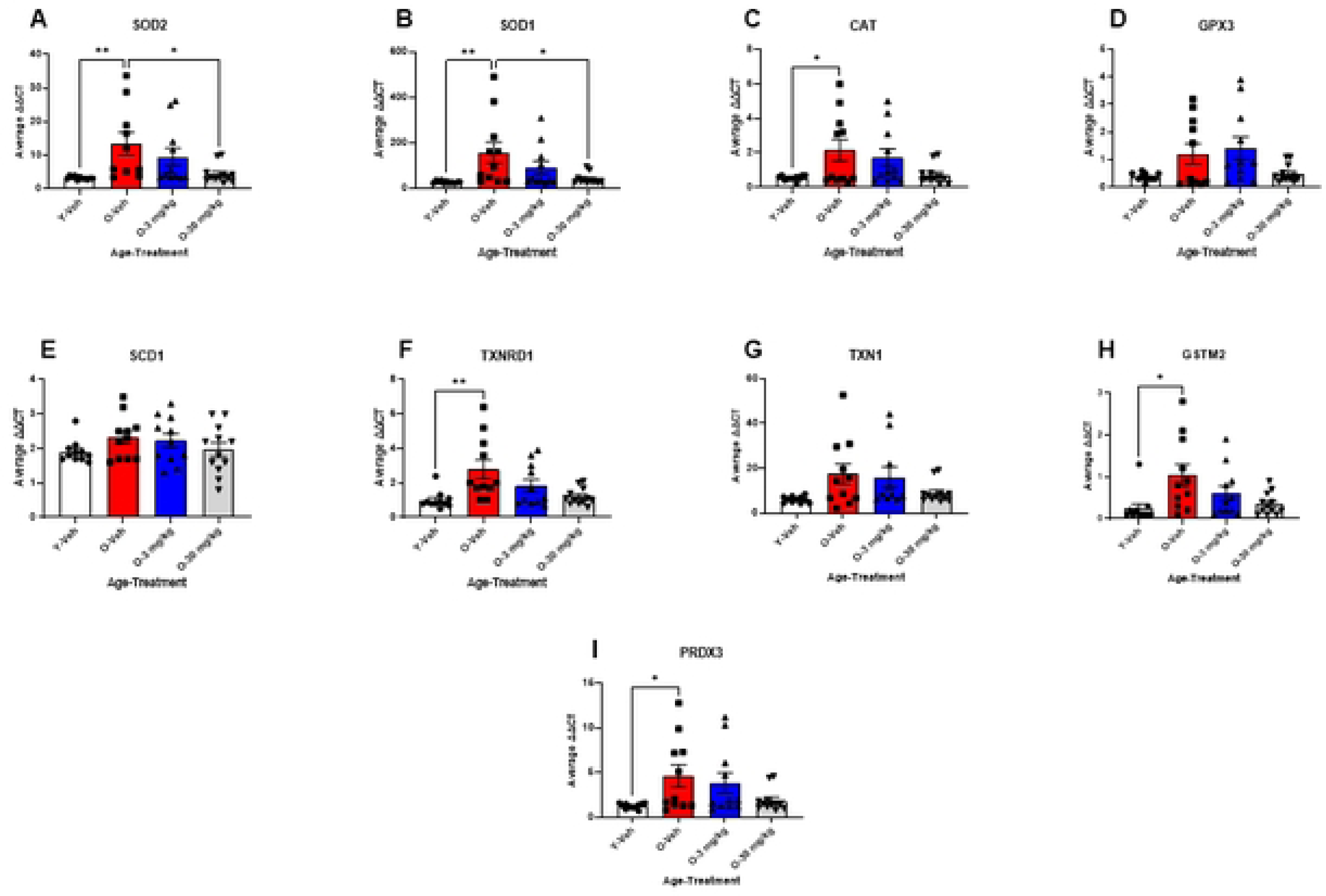
CBD treatment reversed age-related changes in the expression of oxidative stress. Results from qPCR indicate that multiple markers of oxidative stress were elevated with age. 12 mos-old vehicle treated SAMP8 mice (Red bars) had elevated SOD2, SOD1, CAT, TXNRD1 GSTM2, and PRDX3 gene expression (A, B, C, F, H & I) compared to 4 mos-old vehicle treated mice (white bars; A, B, C, F, H & I). Both 3 (blue bars) and 30 mg/Kg (gray bars) CBD treated 12 mos-old SAMP8 mice showed significant decreases in SOD2 and SOD1 gene expression (A & B) with a trend towards a reduction in CAT, TXNRD1, GSTM2 and PRDX3 gene expression (B, F, H, & I). (N = 10∼12 per group for all). * P<0.05.

4-HNE and Gp91phox were measured via western blot to confirm the reduction in oxidative stress. 4-HNE expression was significantly reduced by both 3 mg/Kg and 30 mg/Kg treatment (Figure 4A) and Gp91phox expression was significantly decreased by 30 mg/Kg and no longer significantly elevated by 3 mg/Kg (Figure 4B). Together these findings indicate that CBD treatment reduces oxidative stress in SAMP8 mice.

**Figure 4.**
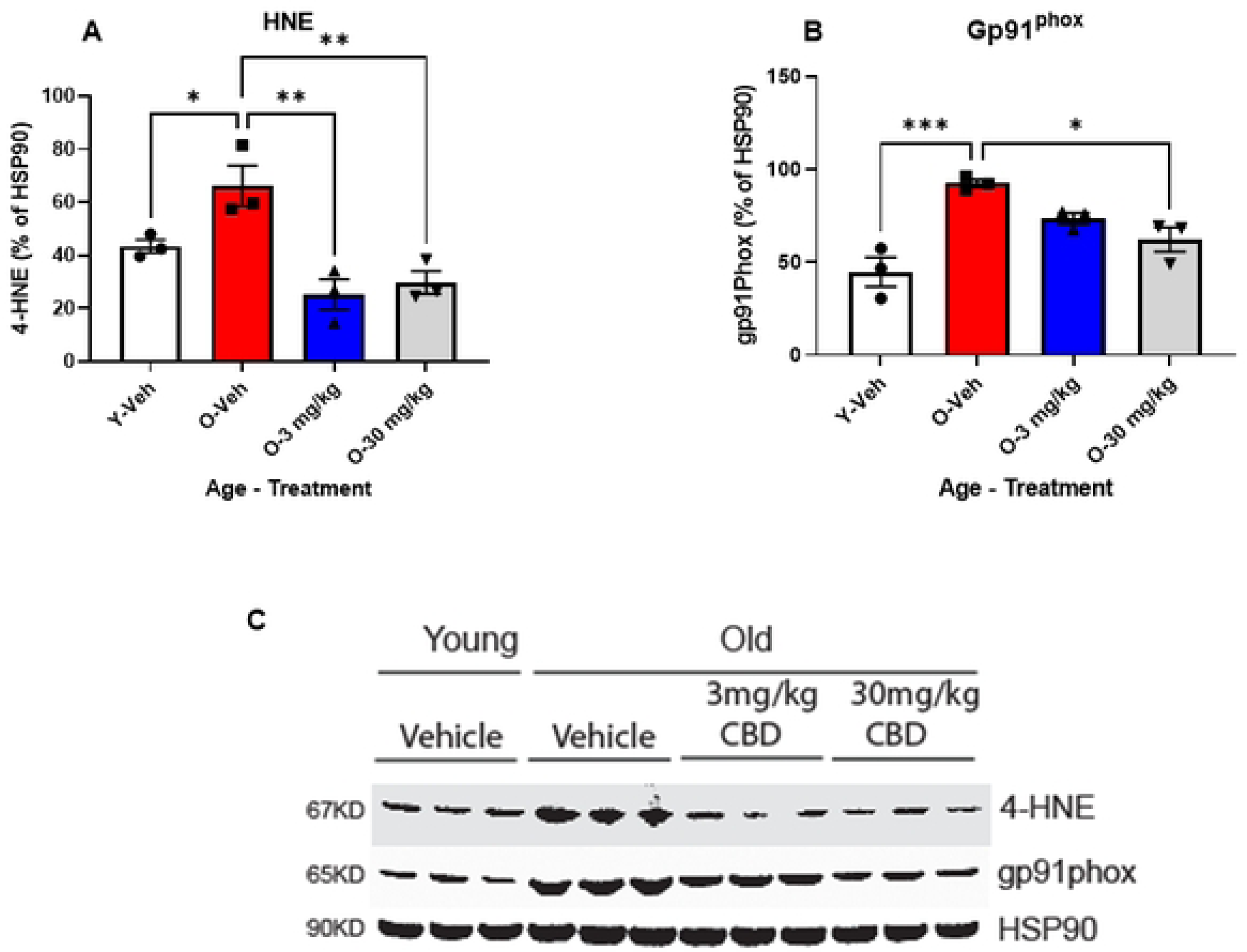
CBD treatment reverses age-related markers of lipid peroxidation and oxidative stress. Western blot analysis of hippocampal tissues taken from 12 mos-old vehicle treated SAMP8 mice (red bars) showed elevated expression of 4-HNE (**A**) and gp91^phox^. (**B**) compared to hippocampal tissues taken from 4 mos-old vehicle treated mice (white bars). Treatment of aged SAMP8 mice with 3 (blue bars) or 30 mg/Kg (gray bars) CBD reduced the expression of both 4-HNE (B) and gp91^phox^ (C) in hippocampal tissues taken from these mice. (N=3 per group for all). C: Depicts a representative Western blot from these experiments. * P<0.05, ** P<0.01, *** P<0.001.

## Discussion

CBD is thought by many to have medicinal properties. Except for a few conditions including two rare forms of epilepsy and multiple-sclerosis-associate spasticity, CBD has not been approved by the FDA (2). There is a need for more preclinical and clinical studies involving CBD. One possible for CBD uses that has been suggested is in the treatment of Alzheimer’s disease. Here we provided daily administration of CBD to male SAMP8 mice. We found that 30 mg/Kg improved learning and memory in both the T maze and NOR. These 2 tasks are considered to have a heavy hippocampal component as demonstrated in work by us and others (30, 31). CBD administration did not change locomotor activity, but it did decrease anxiety in the aged SAMP8 mice. The anxiolytic action of CBD may not be related to aging of the SAMP8 model, as there was no significant difference in between young and old SAMP8 mice treated with vehicle. The aged SAMP8 mice had decreased strength compared to the young mice. CBD did not lead to improvement in measures of strength in the mice. These findings add to a growing body of evidence suggesting that chronic administration of CBD at low doses has a beneficial effect on dementia and anxiety associated with aging (34). In addition, we only used male mice in this study but sex differences have been reported in the pharmacokinetics with greater exposure to the CBD metabolites (35) indicating a study in females is important to assess effectiveness and dose.

Neurodegenerative diseases such as AD, are associated with oxidative stress and inflammation. Postmortem analysis of individuals with AD shows increased markers of oxidative stress such as oxidized proteins, lipid peroxidation, reactive aldehydes and oxidized nucleic acids, in hippocampal and cortical brain regions (36-39). Animal models of AD have shown similar evidence for increased oxidative stress markers in hippocampal and cortical tissues, and investigators have demonstrated a relationship between the oxidative stress and the progression of cognitive impairment in these animals making antioxidant therapies an attractive proposition in AD (40-42). Indeed, treatments with antioxidant properties have also been shown to improve memory in rodent models of AD including the SAMP8 mouse (42-44). However, the results of trials of antioxidants such as Vitamin E (α-tocopherol), C and ubiquinone (CoQ10) as therapies for AD have been disappointing, perhaps due to the large amounts of these antioxidants that are needed to reach effective levels in brain tissue (45, 46).

Our biochemical analysis revealed that SAMP8 mice overall tended to have increases in gene transcripts associated with oxidative stress with aging. However, aged SAMP8 mice treated with 30 mg/Kg CBD had significant reductions in superoxide dismutase 2 (SOD2) and catalase (CAT) transcripts relative to vehicle treated age-matched controls, suggesting a reversal of aging-related changes of oxidative stress. Aged SAMP8 mice also had significantly higher loads of 4-hydroxynonenal (4-HNE), a byproduct of lipid peroxidation, and gp91phox, the catalytic subunit of NADPH oxidase, compared to younger controls. In addition, 30 mg/Kg treated groups had a significant reduction in both 4-HNE in gp91phox. These results are similar to previous results in peripheral tissue which found that CBD reduced 4-HNE and gp91phox in heart and liver (20, 47), and oxidative stress associated with endometriosis (48).

Studies to date on the effects of CBD on dementia have been inconclusive (49). Previous studies involving mouse models of Alzheimer’s disease have found positive effects on cognition. Studies involving AβPPswe/PS1ΔE9 found that 8 months of CBD at 20mg/Kg prevented recognition memory impairment however there was no effect on anxiety, association memory or oxidative damage (50). CBD at 20 mg/Kg was able to prevent spatial memory impairment and cytokine expression induced by Aβ injections in C57BL (51). Esposito *et al.* found that CBD was able to reduce nitric oxidase synthase expression and subsequent nitric oxide production in neurons that had been stimulated with Aβ (52). This same group also found that CBD was able to reduce neuroinflammatory markers IL-1β and inducible NOS expression *in vivo* (53). This was corroborated in the 5xFAD mouse model of familial AD where CBD treatment ameliorated memory impairment through the regulation of IL33 and TREM2 (34). Neuroinflammation, cytokines and oxidative stress are tightly linked contributing to neurodegeneration (54, 55). Here, we found that 30mg/Kg of CBD reduced markers of oxidative stress in aged male SAMP8 mice to levels comparable with young animals.

Our results suggest that 30 mg/Kg CBD treatment improves learning, memory, and reduces anxiety by an oxidative stress modifying mechanism. Given the complex nature of AD pathology and the long incubation period of Aβ and tau effects, targeting broad contributing pathways like oxidative stress or neuroinflammation may provide an effective treatment for those already suffering the symptoms of dementia. Our results also emphasize the need for consideration of the dose. As previously reported in a review by Abate et al. 2021 (34) there is a dose dependent effect with low doses being beneficial and high doses being detrimental.

The use of CBD as a therapy is an exciting development for several reasons. It is easily administered, has a relatively safe profile and can easily cross the blood brain barrier (56-58). The antioxidant effects of CBD have been demonstrated in neurodegenerative cell and animal models (59-61), and although the exact mechanisms by which CBD exerts its antioxidant effects are still under investigation, it appears there are indirect receptor-dependent, as well as direct radical scavenging mechanisms at play (15). The reduction in HNE indicates a decrease in lipid peroxidation. Lipid peroxidation end products play a major role in chronic disease associated with aging including AD (62).

CBD is a relatively weak direct agonist for CB receptors, but it can have a strong indirect effect by inhibiting the metabolism of endogenous endocannabinoids thus allowing for a build-up of their levels within cells. These endogenous cannabinoids, including anandamide, can then activate CB receptors and this may explain some of the neuroprotective effects of CBD that we report here. For example, activation of CB1 receptors on hippocampal neurons is known to protect against the glutamate induced excitotoxicity, that has been hypothesized as a mechanism of neurodegeneration in AD (63, 64). Furthermore, indirect activation of CB2 receptors on microglia and astrocytes can directly decrease the production of reactive oxygen species (ROS) and other inflammatory factors from enzymatic systems in those cells (65-67). This latter effect fits well with our observation of the prevention of the upregulation of GP91^phox^ (a subunit of the ROS producing enzyme NADPH oxidase) in hippocampal preparations of SAMP8 mice treated with CBD. Our results also fit well with other studies examining the effect of CBD and related compounds on NADPH oxidase expression (68, 69).

Another way that CBD could be neuroprotective in our studies lies in its ability to directly scavenge ROS thus preventing their reactions with lipids, proteins and DNA (20). This antioxidant scavenging action of CBD has been observed in numerous cellular and neuronal models including those challenged with Aβ, and its neuroprotective effect has been shown to be more potent than antioxidants such as Vitamins E or C (20, 52, 70, 71). This would certainly contribute to our findings that expression of 4-HNE is prevented in SAMP8 mice treated with CBD. It is also unsurprising that our results show a decrease in the expression of antioxidant enzymes such as SOD and Catalase, as well as constituents of glutathione production with CBD treatment. These antioxidant systems are upregulated in response to oxidative stress such as that evident in our untreated SAMP8 mice, and the oxidant levels in CBD treated SAMP8 mice are too low to require the upregulation of these antioxidant defenses.

Our results indicate that CBD can reverse age-related changes in the SAMP8 mice. CBD reversed memory impairment in both the T-maze and NOR with a 24 hour retention interval. These results indicate that CBD is able to reverse memory impairment in both spatial and recognition tasks (30, 31) Taken together therefore, there appears to be an antioxidant effect of CBD contributing to the improvement in memory in the aged mice. These results show that CBD is an attractive therapeutic warranting further investigation in AD, and other neurodegenerative diseases.

## Author Contributions

MNG contributed to study design, performed experiments, analyzed data, and wrote first draft; SB, BJY and MLN performed experiments and analyzed data; HM, AAB, and JEM contributed to study concept and edited the manuscript; SAF designed the study, oversaw experiments and data analysis, and edited the manuscript. All authors read and approved the final manuscript.

## Acknowledgements

This work was supported by the Doisy Research Fund, St Louis University School of Medicine. The CBD was donated by BeLeaf.

